# From *mec* cassette to *rdhA*: a key *Dehalobacter* genomic neighborhood in a chloroform and dichloromethane–transforming microbial consortium

**DOI:** 10.1101/2023.11.10.566631

**Authors:** Olivia Bulka, Katherine Picott, Radhakrishnan Mahadevan, Elizabeth A. Edwards

## Abstract

Chloroform (CF) and dichloromethane (DCM) are groundwater contaminants of concern due to their high toxicity and inhibition of important biogeochemical processes such as methanogenesis. Anaerobic biotransformation of CF and DCM has been well documented but typically independently of one another. CF is the electron acceptor for certain organohalide-respiring bacteria that use reductive dehalogenases (RDases) to dechlorinate CF to DCM. In contrast, known DCM-degraders use DCM as their electron donor, which is oxidized using a series of methyltransferases and associated proteins encoded by the *mec* cassette to facilitate the entry of DCM to the Wood-Ljungdahl pathway. The SC05 culture is an enrichment culture sold commercially for bioaugmentation, that transforms CF via DCM to CO_2_. This culture has the unique ability to dechlorinate CF to DCM using electron equivalents provided by the oxidation of DCM to CO_2_. Here we use metagenomic and metaproteomic analysis to identify the functional genes involved in each of these transformations. Though 91 metagenome-assembled genomes were assembled, the genes for an RDase—named *acdA*—and a complete *mec* cassette were found to be encoded on a single contig belonging to *Dehalobacter*. AcdA and critical Mec proteins were also highly expressed by the culture. Heterologously-expressed AcdA dechlorinated CF and other chloroalkanes but had 100-fold lower activity on DCM. Overall, the high expression of Mec proteins and the activity of AcdA suggest a *Dehalobacter* capable of dechlorination of CF to DCM, and subsequent mineralization of DCM using the *mec* cassette.

**Importance:** Chloroform (CF) and dichloromethane (DCM) are regulated groundwater contaminants. A cost-effective approach to remove these pollutants from contaminated groundwater is to employ microbes that transform CF and DCM as part of their metabolism, thus depleting the contamination as the microbes continue to grow. In this work, we investigate bioaugmentation culture SC05, a mixed microbial consortium that effectively and simultaneously degrades both CF and DCM coupled to the growth of *Dehalobacter*. We identified the functional genes responsible for the transformation of CF and DCM in SC05. These genetic biomarkers provide a means to monitor the remediation process in the field.

## Introduction

Chlorinated alkanes like chloroform (CF) and dichloromethane (DCM) are prevalent and persistent groundwater contaminants. They have been widely used in many industrial processes as solvents and degreasers, resulting in serious groundwater contamination (1, 2). CF is also generated as a by-product of water disinfection and by the dechlorination of carbon tetrachloride (1, 3). CF and DCM are both formed naturally (4), but at much lower concentrations than those resulting from anthropogenic sources. Both CF and DCM are toxic and carcinogenic to humans (5, 6), but CF causes additional environmental problems by inhibiting natural biogeochemical cycling and dechlorination of chloroethenes at mixed-contaminant sites (1, 7–9). Through biostimulation and bioaugmentation, mixed microbial cultures—often derived from contaminated sites—are an effective method to remove these compounds from anaerobic contaminant sites (Sandra Dworatzek, personal communication, 1, 2, 8).

Numerous anaerobic enrichment cultures and isolated organisms targeting CF or DCM have been described, but typically these cultures focus solely on either CF or DCM. CF-reducing enrichment cultures typically accumulate DCM as a product of CF dehalogenation, without degradation of DCM (10–14). Moreover, cultures enriched on DCM can be sensitive to CF concentrations, which leads to difficulties with sequential remediation (15). For example, when CF-dechlorinating culture ACT-3 was co-cultured with DCM-degrading Consortium RM, DCM mineralization only occurred 100-200 days after CF was completely transformed to DCM (16). Despite the structural similarity of CF and DCM, their known mechanisms of biotransformation are distinct.

All known anaerobic CF-dechlorinating organisms are classified as *Dehalobacter* spp. (10–12, 17), with the exception of *Desulfitobacterium* sp. strain PR (14). All *Dehalobacter* and many *Desulfitobacterium* strains are organohalide-respiring bacteria (OHRB)—organisms that use organohalides as terminal electron acceptors in their respiratory processes. These strains efficiently respire CF through reductive dechlorination, yielding DCM as a product (10–12, 14, 17). These reactions are catalyzed by enzymes called reductive dehalogenases (RDases). RDases are ubiquitous among OHRB, and each of the known CF-respiring organisms expresses a slightly different active RDase within the same classification group.

To classify characterized RDases and uncharacterized reductive dehalogenase homologous genes (*rdhA*) from (meta)genome sequences, all predicted proteins have been organized into ortholog groups (OGs) (18, 19). Each OG comprises protein sequences sharing more than 90% amino acid sequence identity (19). All known CF-dechlorinating organisms possess RDases that cluster in a single group: OG 97 (similarity matrix in Figure S1). It is worth noting that despite having highly similar amino acid sequences, the OG 97 RDases are distinct in their substrate profiles, which include various structurally related halomethanes and chloroethanes. Members of OG 97 that are capable of CF dechlorination include CfrA from *Dehalobacter* sp. CF (20), TmrA from *Dehalobacter* sp. UNSWDHB (21), ThmA from *Dehalobacter* sp. THM1 (12), CtrA from *Desulfitobacterium* sp. PR (14), and most recently RdhA D8M_v2_40029 from *Dehalobacter* sp. 8M (22). Only *Desulfitobacterium* sp. PR—and by inference, its enzyme CtrA—has been documented to dechlorinate DCM to chloromethane (CM), but it is a minor transformation product (5% of DCM converted to CM) and DCM has not been shown to sustain growth of this organism (14).

The four known anaerobic DCM-degrading microorganisms use DCM as electron donor in one of two established anaerobic pathways. Both pathways harness the Wood-Ljungdahl Pathway (WLP) and involve the *mec* cassette in the initial DCM assimilation step (23). This 10-gene cassette encodes a methyltransferase system that is thought to assimilate DCM to methylenetetrahydrofolate to launch both pathways before they diverge later in the WLP (23). The DCM fermentation pathway—first discovered in *Dehalobacterium formicoaceticum* strain DMC—uses the WLP to release chloride ions from DCM, ultimately producing formate and acetate (24, 25). *Candidatus* Formimonas warabiya and *D. formicoaceticum* strain EZ94 are also predicted to use this pathway (26– 28). The second pathway, DCM mineralization, also employs the WLP to metabolize DCM, but instead reduces carbon dioxide and hydrogen as end products (29). This mechanism has only been shown in *Candidatus* Dichloromethanomonas elyunquensis, which was originally classified as a strain of *Dehalobacter* when first identified using its 16S rRNA sequence alone (29–31).

Other studies have implicated *Dehalobacter* strains in DCM degradation, however none has been genomically verified. A *Dehalobacter* strain was reported to be detected in consortium CFH2, which later became dominated by *Ca*. F. warabiya during subsequent enrichment (26, 32). This *Dehalobacter* strain disappeared from the culture before genomic analysis could confirm its predicted taxonomy (26). In our previous work (33), we identified a *Dehalobacter* strain that persisted in DCM-amended subcultures of the SC05 consortium. In this study, we continued to investigate the role of *Dehalobacter* in DCM metabolism in the SC05 culture.

SC05 is a microbial consortium maintained by SiREM (Guelph, ON) that dechlorinates CF to DCM and concurrently mineralizes DCM to carbon dioxide and hydrogen under anaerobic conditions (34). We previously established a subculture at the University of Toronto, known as SC05-UT, which has been maintained continuously on CF alone, without exogenous electron donor addition, for >1000 days (33). This culture produces sufficient hydrogen through DCM mineralization (2 mol H_2_ produced from 1 mol DCM) to power its CF dechlorination (1 mol H_2_ needed for reduction of 1 mol CF) (33). From this culture, an additional sub-transfer was made and maintained on DCM alone, which was named DCME (33). In each of these cultures, the same *Dehalobacter* 16S amplicon sequence variant (ASV) increased in abundance while the culture consumed CF and DCM, as well as DCM alone (33). These findings suggested a multifunctional *Dehalobacter* that may produce its own electron donor from mineralization of DCM (its dechlorination product), but the mechanisms for both CF dechlorination and DCM mineralization have remained uncharacterized in this culture. Here we searched for functional genes in metagenomes from SC05-UT and DCME to identify and characterize the RDase dechlorinating CF, to provide insight into the mechanism of DCM degradation in the SC05-UT and DCME cultures, and finally to determine if the genes encoding these pathways belong to *Dehalobacter*.

## Results & Discussion

### 1. Dehalobacter metagenome-assembled genomes

A more detailed description of the metagenome assemblies and bins in this study can be found in our related resource announcement (35), and overall metagenome quality statistics can be found in Table S5. Briefly, forty draft metagenome-assembled genomes (MAGs) were assembled from the DCME metagenome, and 51 draft MAGs were assembled from the SC05-UT metagenome, representing 53% and 77% of total microorganisms, respectively, as determined by relative abundance estimates in Anvi’o (36). One *Dehalobacter* MAG was assembled from each metagenome. In the SC05-UT metagenome, the *Dehalobacter* MAG was 3.46 Mb in 7 contigs and 98.6% complete as estimated by Anvi’o (N_50_ = 572 kb, mean coverage = 1028, read recruitment = 12.3%) (Table S1). In DCME, the MAG was only 1.97 Mb in 28 contigs (N_50_ = 148 kb) and estimated as 90.1% complete (Table S1), though lacking more than 1 Mb compared to other closed *Dehalobacter* genomes. Mean read coverage (42.3) and read recruitment (0.3%) were lower for the DCME *Dehalobacter* MAG than that from SC05-UT (Table S1). Read recruitment is defined as reads mapped to a bin divided by total reads in the metagenome, which is skewed by the PCR amplification of the metagenomic reads. The abundances determined by amplicon sequencing are more reliable, which show the *Dehalobacter* abundance is 10% of total bacteria at the time of sequencing.

The DCME metagenome was sequenced after the subculture had been enriched on DCM only for more than one year, during which many methanogens flourished, as well as other microorganisms whose growth are typically inhibited by CF (Figure S2). As a result, *Dehalobacter* had a lower relative abundance in the DCME culture compared to SC05-UT at the time of sequencing (10-20% vs. 70-90% as determined by 16S amplicon sequencing, Figure S2), which led to a much more fragmented and incomplete *Dehalobacter* MAG with lower read coverage overall compared to the *Dehalobacter* MAG from SC05-UT. Despite this poor quality, our previous work has shown that only *Dehalobacter* grows as DCM is degraded, which indicates it is in fact a DCM-degrader despite its lower abundance than other MAGs. Other draft MAGs are summarized in Table S1.

### 2. Identification of AcdA, the expressed RDase in SC05-UT

To identify the CF-dechlorinating RDase in SC05-UT, its metagenome was first assembled and searched for *rdhA* genes. Of the 37 identified *rdhA* genes, 27 were from contigs in the *Dehalobacter* MAG (Table S2). Closed *Dehalobacter* genomes typically encode ∼20-25 RDases, but each strain often only expresses one for respiration (19, 37, 38). Of the remaining 10 *rdhA* sequences, three were on unbinned contigs, while seven were binned as Anaerolineaceae, Mesotoga, or Synergistales (Table S2).

One RDase was identified in the proteome of SCO5-UT with a high protein abundance ranking (ID: JAWDGN_38927, mean protein abundance rank: 3; Table S3A). While not quantitative, a high protein abundance ranking (with 1 as the highest rank) implies a greater expression level of that protein than those with a lower rank. A highly similar RDase with five amino acid differences was also the only to be expressed in DCME (ID: JAWDGO_25608, mean protein abundance rank: 4; Table S3B). Like other characterized *rdhA* sequences, this gene contains a twin-arginine transport (TAT) sequence and two iron-sulfur cluster binding motifs. Immediately downstream are an *rdhB* and an *rdhC* gene, between which is an inserted transposase gene, before a terminator sequence to end the operon (Figure S3, Table S6). In the DCME metagenome, no transposase interrupts the *rdhB* and *rdhC* genes.

This expressed SC05-UT RDase was previously studied using compound-specific isotope analysis (CSIA)—an analytical technique used to provide insight into transformation pathways—wherein the RDase was referred to as KB-1^®^ Plus CF RdhA (39). No biochemical analysis was performed at that time. The RDase clustered with OG 97, the ortholog group containing all five known CF-dechlorinating RDases, with its highest amino acid similarity (96.7%) to TmrA (WP_034377773) (39). CSIA has revealed distinct fractionation patterns for CF dechlorination, indicating differing mechanisms of action or rate-limiting steps for four mixed cultures expressing the RDases, TmrA, CfrA, RdhA D8M_v2_40029, and the SC05-UT RDase (17, 39, 40). A handful of amino acid variants within the active sites were speculated to cause these fractionation differences (39). The isotope fractionation pattern differentiated the SC05-UT RDase from the other CF-dechlorinating RDases, suggesting that it is another distinct RDase within OG 97 (39). We have observed that it is indeed the case that these active site residue differences have an influential effect on RDase substrate preferences (41; manuscript in preparation). The OG 97 classification, in addition to its lone expression from the metagenome’s roster of putative RDases, identifies AcdA as a chloroform reductive dehalogenase.

### 3. AcdA dechlorinates many chloroalkanes, but not DCM

To explore AcdA’s dechlorinating activity, we expressed the native gene heterologously in *Escherichia coli*. Its expression was much lower than what has been previously seen for TmrA using the same method (42). This could be because the native *Dehalobacter* gene was not codon-optimized for expression, or because the enzyme itself might be less soluble. Regardless, the enzyme was enriched by nickel-affinity chromatography, and the protein concentration used in assays was adjusted using the estimated purity of 20%. The expressed AcdA was assayed against substrates accepted by other RDases in OG 97 including CF, 1,1,1-trichloroethane (TCA), 1,1,2-TCA, and 1,1-dichloroethane (DCA) (Figure 1). DCM was also tested as a substrate to investigate if there was sufficient activity to contribute to DCM mineralization (Figure 1A). Enzyme-free negative controls were used for each substrate. No product was detected in the controls, except for one CF replicate (<0.5 nmol DCM) and one 1,1-DCA replicate (<0.5 nmol vinyl chloride [VC]) (Supplemental Information, Dataset S4). TmrA was used as a positive control and as a comparison against a known CF reductase. The enzyme activity is reported here as nmol of product measured per second of the reaction per mg of RdhA enzyme (enzyme free control is in nmol/s).

**FIGURE 1.**
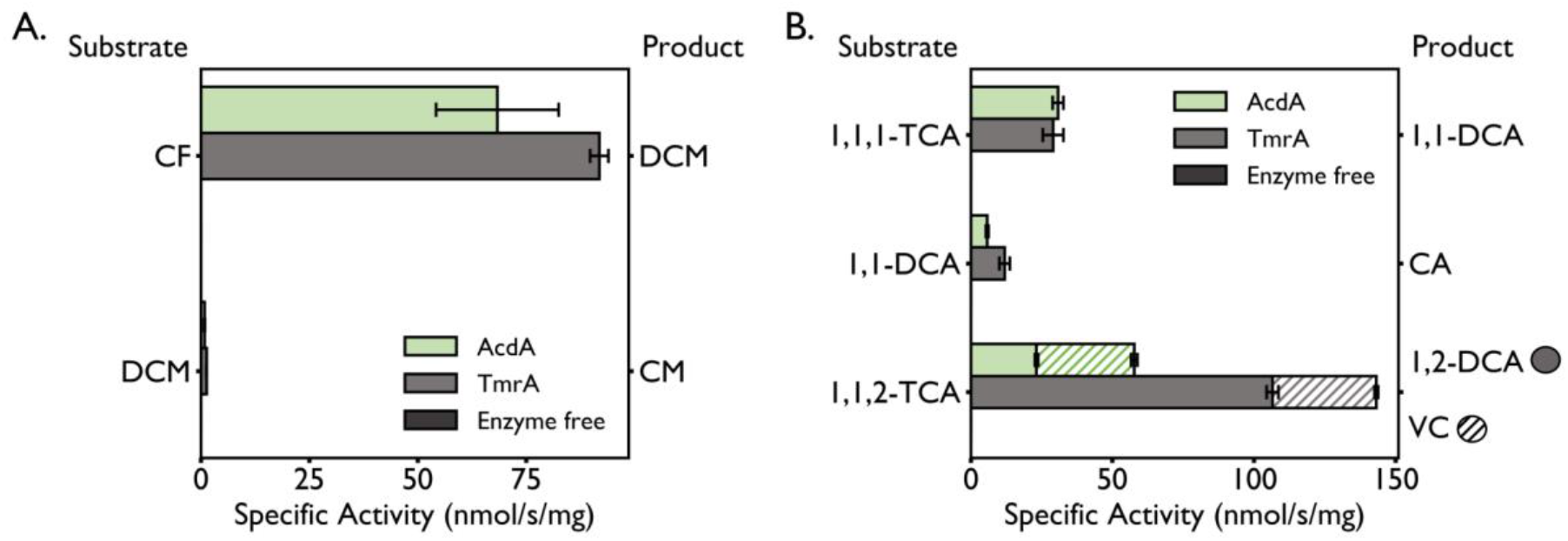
AcdA and TmrA specific activity on chlorinated A) methanes and B) ethanes. Specific activity is the amount of product (nmol) per second per mg of RDase in the reaction, enzyme free negative control is in nmol/s. Reactions were incubated at room temperature for 1 h with no headspace. Activity on 1,1,2-TCA is stacked depending on the product of the reaction: 1,2-DCA (solid) or VC (striped). Error bars are the standard deviation between triplicate reactions. CF = chloroform; DCM = dichloromethane; CM = chloromethane; TCA = trichloroethane; DCA = dichloroethane; CA = chloroethane; VC = vinyl chloride.

Generally, AcdA displayed comparable activity levels on all tested substrates to TmrA (Figure 1). Similar to TmrA and other OG 97 enzymes, AcdA sequentially dechlorinates 1,1,1-TCA to 1,1-DCA, and then 1,1-DCA to chloroethane (CA), though it dechlorinates 1,1-DCA at a much slower rate. Interestingly, the enzyme assay was able to predict 1,1,1-TCA and 1,1-DCA as putative growth substrates for SC05-UT. We confirmed that the culture could dechlorinate 1,1,1-TCA and 1,1-DCA, albeit ∼10 times more slowly than CF, with CA accumulating as a final dechlorination product (Text S5, Figure S4). AcdA also dechlorinated 1,1,2-TCA via two different reaction pathways yielding a mixture of 1,2-DCA and VC at a ratio of 2:3. This is a much higher ratio of VC than most OG 97 RDases produce, and it may be of interest to investigate the cause of this mechanistic switch.

AcdA is most active on CF as a substrate, which is unsurprising given that the SC05-UT culture originated from a CF contamination site (34). While there is a small amount of DCM dechlorination observed (0.79 ± 0.08 nmol/s/mg), this activity is 100-fold lower than that of CF dechlorination (68 ± 14 nmol/s/mg). CtrA and TmrA also display nominal dechlorination activity on DCM (14; Figure 1A). DCM dechlorination is predicted to be co-metabolic in *Desulfitobacterium* sp. PR, whereas no DCM metabolism has been observed in *Dehalobacter* sp. UNSWDHB (11, 14). Given AcdA’s low activity on DCM compared to CF, and the sustained DCM removal from the SC05-UT culture (33), AcdA is not likely to be the primary DCM-degrading enzyme.

### 4. Only the *Dehalobacter* MAG contains the *mec* cassette

To predict a mechanism for DCM degradation in SC05, we searched both the SC05-UT and DCME metagenomes for the *mec* cassette: a ten-gene cassette identified specifically in the genomes of all known DCM-degrading microorganisms (23). Only one instance of the *mec* cassette was found in each metagenome—on contigs assigned to each *Dehalobacter* MAG (SC05-UT contig: JAWDGN010000066, 364.8 kb, mean coverage: 1561.9; DCME contig: JAWDGO010000051, 279.2 kb, mean coverage: 37.7). The more fragmented assembly of the DCME *Dehalobacter* MAG resulted in a broken DCME *mec* contig that terminates after the *mecH* gene (Figure 2). A search for the subsequent *mecIJ* genes in the other contigs from the DCME metagenome revealed one contig in an Anaerovoracaceae bin (contig JAWDGO010000514, 53.6 kb). This contig had markedly higher GC content than the rest of the gene neighborhood in both *Dehalobacter* MAGs (47.1% vs. 40.3–40.8%). Moreover, no long-read coverage linked *mecGHIJ* to the rest of the contig, indicating a potential misassembly (Figure S5). For these reasons, this contig was not merged with the *Dehalobacter mec* contig JAWDGO010000051 (expanded upon in Supplementary Text S6). An alternative downstream sequence encoding a transposase and *mecIJ* was resolved using PacBio HiFi reads to perform contig extension on the end of contig JAWDGO010000051, which is syntenic to the *mec* cassette seen in the SC05-UT *Dehalobacter* MAG and share 100% nucleotide identity (Supplementary Text S6, Figure S7). Future sequencing efforts will attempt to connect the 3’ end of the *mec* cassette to the *Dehalobacter* genome in the DCME culture.

**FIGURE 2.**
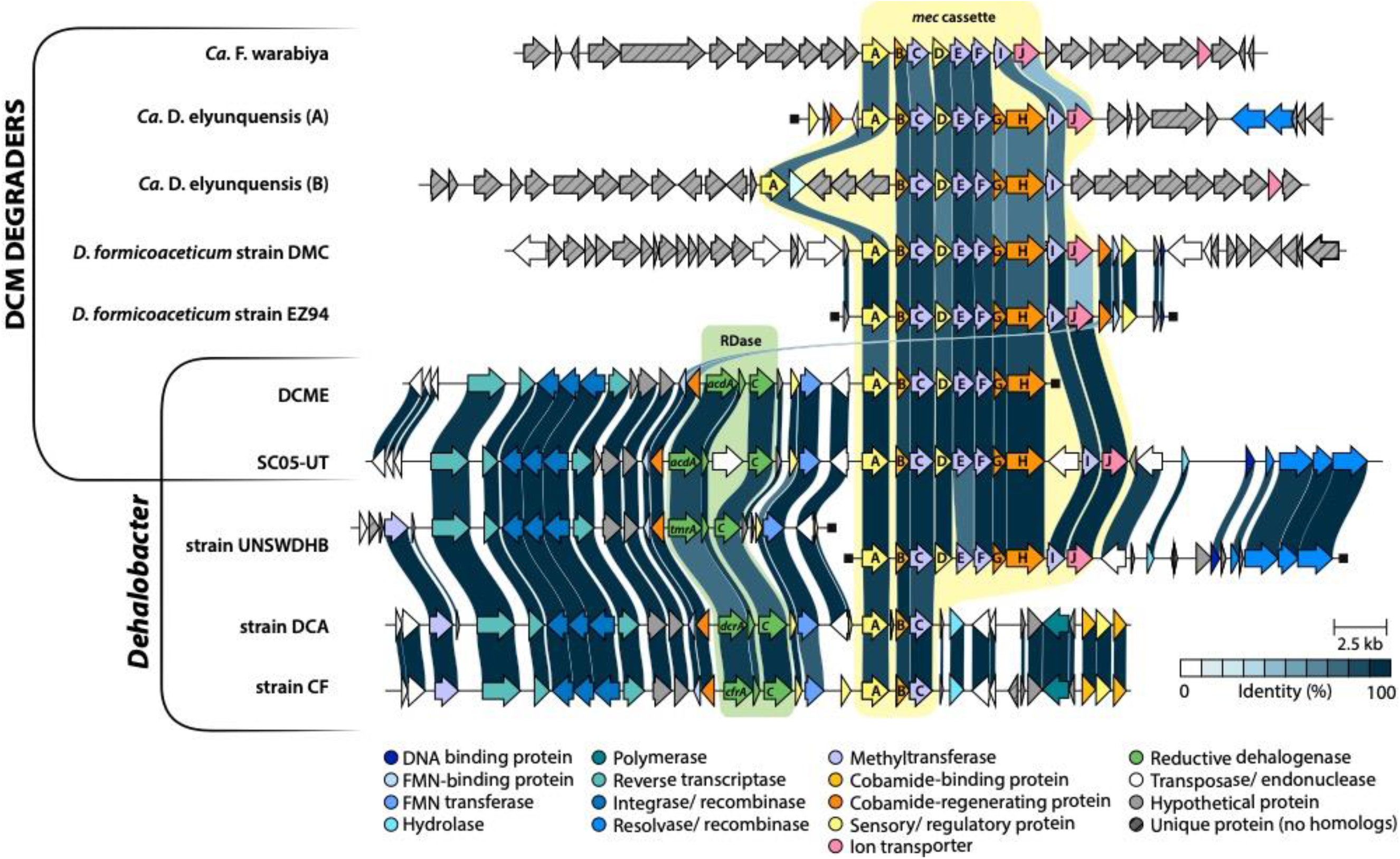
Genomic neighbourhoods of all known *mec* cassettes in DCM degrading organisms and *Dehalobacter* strains. RDase genes are highlighted in green; *mec* genes are highlighted in yellow. Homologous genes are color-coded by function; links between homologs are colored by percent nucleotide identity. Genes with no local homologs are grey striped. Ends of a contig are capped with black squares.

One bin in the DCME metagenome was assigned to Dehalobacteriia—the class containing Dehalobacteriales, which includes the DCM degrading *Dehalobacterium* spp. and *Ca*. F. warabiya, however, this bin did not contain a *mec* cassette (Table S1). The Dehalobacteriia class contains two orders other than Dehalobacteriales: UBA7702 and UBA4068, neither of which include microbes known to degrade DCM nor contain the machinery to do so (43). The DCME Dehalobacteriia MAG is of the order UBA7702, one of the two non-DCM degrading orders, so is more likely an amino acid scavenger in the culture, like others in this class of organisms (43). Moreover, this bin was assembled in the DCME culture metagenome, but was not identified in the SC05-UT culture metagenome (Table S1). Accordingly, the relative abundance of the Dehalobacteriia strain using 16S rRNA gene amplicon sequencing at the time of metagenome sequencing was between 2-22% abundance in DCME but was at < 0.4% abundance in SC05-UT (Figure S2). At such low abundance in the parent culture, and lacking the *mec* cassette, we do not expect this organism to be the DCM degrader in either culture. The growth of this microbe may have been inhibited by CF, as many fermenters are, and thus proliferated during DCM enrichment due to the removal of the CF pressure.

### 5. One gene neighborhood encodes *acdA* and the *mec* cassette

In both the SC05-UT and DCME metagenomes, *acdABC* is encoded on the same contig as the *mec* cassette, less than 4 kb upstream of *mec*A (Figure 2), further supporting the cassette’s assigned *Dehalobacter* taxonomy. When aligned, most genes in this region share 100% nucleotide identity. Exceptions include variability in *acdA* (99.6% identity) and *acdC* (97% identity). Though *mecB* and *mecC* also differed between the initial assemblies of the two MAGs (93% and 95% nucleotide identity, respectively), mapping of PacBio HiFi reads could correct the short-read mapping ambiguities that led to these variations. There are also 10 additional single nucleotide polymorphisms throughout the region, and a transposase insertion in SC05-UT, both of which may be indicative of population variation between (and within) the cultures.

A gene neighbourhood including ∼10 kb up and downstream of the RDase and *mec* cassette was extracted from published genomes of all known DCM-degraders and *Dehalobacter* strains known to possess a full or partial *mec* cassette. The SC05-UT *mec* cassette is syntenic to the other DCM-degraders, except for a transposase inserted between *mecH* and *mecI* (Figure 2). The SC05-UT cassette is most similar to that of *Dehalobacter* sp. UNSWDHB (Figure 2), which lacks the ability to degrade DCM despite encoding a “full” cassette of all 10 *mec* genes (11). This is speculated to be due to its truncated *mecE*, which encodes a critical methyltransferase for assimilation of DCM into the WLP (23). Two other strains of *Dehalobacter* (sp. CF and sp. DCA) also contain three genes homologous to *mecABC*, but cannot degrade DCM for lack of the remaining *mec* genes (10, 37).

Interestingly, each of these *Dehalobacter* strains has a characterized OG 97 RDase upstream of *mecA*, besides *Dehalobacter* strain UNSWDHB, whose *mec* contig ends just upstream of the *mecA*. An additional contig from strain UNSWDHB was included in the gene neighborhood comparison to include its OG 97 RDase, *tmrA*. These *Dehalobacter* strains, including strain UNSWDHB, share a syntenic region of at least 10 homologous genes upstream of the *rdhA—*including reverse transcriptase and integrase genes, hypothetical proteins, and cobamide and FMN binding proteins—and at least two genes between the *rdhC* and *mecA*—homologous regulatory and FMN transfer proteins (Figure 2). To join the two contigs from strain UNSWDHB, only 117 bp are missing from the assembly if the contigs are indeed adjacent and syntenic with the other *Dehalobacter* strains. This may be resolved by further sequencing or assembly efforts.

Downstream of the *mec* cassette, the *Dehalobacter* strains show two distinct gene signatures. Strains CF and DCA, which encode only *mecABC*, also encode different downstream genes (hydrolases, cobamide-binding proteins) than the contigs from strain UNSWDHB and the SC05-UT MAG (recombinases), which share homology. A rearrangement may have occurred in this region in strains CF and DCA, resulting in a loss of *mecDEFGHIJ*. Strains CF and DCA were originally enriched from a site contaminated with 1,1,1-TCA (44), so the lack of selective pressure to maintain DCM degradation machinery may have facilitated this loss. This genomic area is also riddled with transposases, implying an area of high genomic change, which may also play a part in gene loss and acquisition.

Despite this precedent for partial, nonfunctional, *mec* cassettes in CF-dechlorinating *Dehalobacter* strains, the SC05-UT *mec* cassette is the first reported to be found in full—and evidently functioning—in a *Dehalobacter* genome. Enzyme assays for characterizing the Mec cassette must be developed before the function of this pathway can truly be confirmed in SC05-UT or any other DCM-degrader. In the meantime, expression levels of these proteins during active degradation do serve as a proxy for their importance in the culture, as described below.

### 6. Expression of DCM degradation genes

To compare the expression of the Mec proteins in SC05-UT (while CF and DCM were degraded) and DCME (while DCM alone was degraded), a proteomic workflow was performed on each culture. The annotations used for proteomic searching of each metagenome are provided in the supplementary information, Datasets S1 and S2. All detected peptides and proteins are shown in the supplemental information, Dataset S3, and all protein hits mapping to *Dehalobacter* are filtered in Table S3. Most proteins encoded by the *mec* cassette are expressed under both culture conditions (Figure 3); the most highly expressed are MecE, MecF, MecB and MecC, which each rank in the top ten proteins expressed in at least one culture. The high proteomic rank of these proteins and the proximity of several strong promoters on the contig suggest a region of high expression in the genome (Figure S3).

**FIGURE 3.**
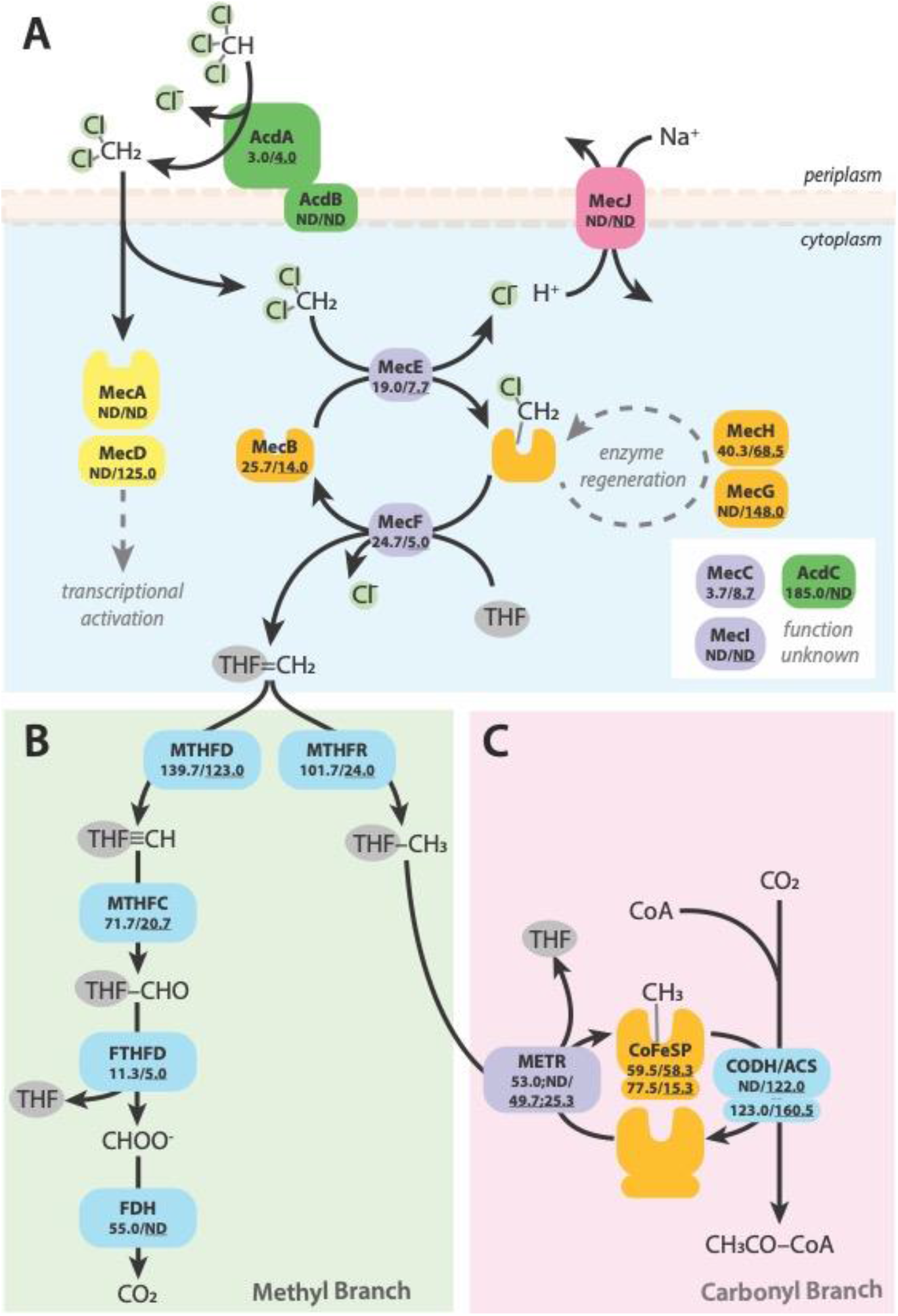
Protein expression of key pathways in SC05-UT and DCME, including A) CF and DCM metabolism, and B) methyl and C) carbonyl branches of the Wood-Ljungdahl pathway. Average rank of biological replicates (n = 3) displayed as SC05-UT/<u>DCME</u>, where high rank (a low number) suggests high abundance. ND: not detected by proteomics, and expression of multi-copy genes are separated with “;”. THF: tetrahydrofolate, MTHFD: methylene-THF dehydrogenase, MTHFC: methenyl-THF cyclase, FTHFD: formyl-THF deformylase, FDH: formate dehydrogenase, MTHFR: methylene-THF reductase, METR: 5-methyl-THF:corrinoid/iron-sulfur protein methyltransferase, CoFeSP: corrinoid/iron-sulfur protein, CODH/ACS: CO dehydrogenase/acetyl-CoA synthase complex. Locus tags and enzyme descriptions further described in Table S4.

The only Mec proteins not detected in either subculture were MecA, MecI, and MecJ. Notably, MecI and MecJ are located after the inserted transposase found in the cassette which may interrupt their expression (Figure 2). A terminator sequence is predicted to exist within the transposase gene, which could account for this effect (Figure S3). Additionally, MecA was not detected in *D. formicoaceticum* strain EZ94, and MecJ has never been detected in studies of either *D. formicoaceticum* strain or *Ca*. Dichloromethanomonas elyunquensis (23, 28). Overall, high expression of the key methyltransferases and corrinoid proteins show the possibility of DCM assimilation to methylenetetrahydrofolate in SC05, after which methylenetetrahydrofolate could then enter the WLP for further metabolism (Figure 3).

We also searched the metagenomes and metaproteomes for genes representing each step of the WLP, all of which were found encoded in each *Dehalobacter* MAG, after the inclusion of one unbinned contig annotated as belonging to *Dehalobacter* by MetaErg (JAWDGO010000155) in the DCME *Dehalobacter* MAG (Table S4). The WLP is a critical pathway for carbon assimilation and energy conservation in many OHRB (37, 45, 46). This pathway becomes doubly crucial when the *mec* cassette is in use, as it must also accept the methylenetetrahydrofolate produced from DCM assimilation and convert it to carbon dioxide or acetate.

In SC05-UT, while degrading CF and DCM, as well as in DCME while degrading DCM alone, this *Dehalobacter* population expresses almost all WLP enzymes (Figure 3, Table S4). Two exceptions are formate dehydrogenase (FDH, Figure 3B) in DCME, and the alpha subunit of the carbon monoxide dehydrogenase/ acetyl-CoA synthase complex (CODH/ ACS, Figure 3C) in SC05-UT. It is possible that expression of these two proteins was low enough to evade detection in our proteomics measurements, or that a more highly-expressed version of this gene with a divergent sequence is missing from the *Dehalobacter* MAG or the metagenome and therefore not detected in our protein search. In the case of SC05-UT, acetate-CoA ligase was expressed, through which acetyl-CoA could be produced from exogenous acetate without CODH/ ACS (Table S4). This suggests that an active acetogen in the culture is providing *Dehalobacter* with a pool of acetate in SC05-UT that may not be readily available in DCME. Nonetheless, the expression of the WLP and the Mec proteins positions the *mec* cassette as a likely mechanism of DCM assimilation in SC05.

## Conclusions & Implications

SC05-UT simultaneously dechlorinates CF to DCM and mineralizes DCM to carbon dioxide, and its *Dehalobacter* population encodes both an active RDase and the complete *mec* cassette. Here we introduce AcdA—another chloroform dechlorinating RDase in OG 97 that expands the arsenal of biochemical tools against chloroalkane contaminants. In addition to AcdA, the Mec cassette and WLP proteins were among the most highly expressed in the metaproteome, indicating likely mechanisms for CF and DCM metabolism by a *Dehalobacter* population.

One contig contains both *acdA* and the *mec* cassette, concluding one organism is genomically capable of both biotransformation steps. A single *Dehalobacter* strain may truly perform both metabolisms simultaneously, or subtle strain differences may exist and distinguish two metabolically different *Dehalobacter* strains. Though obtaining one or more *Dehalobacter* isolates would provide definitive answers to this question, these organisms rely on a complex microbial community to provide the correct balance of nutrients to support their growth. This restricted metabolism in addition to long doubling times pose challenges in obtaining a pure culture, though isolations are not impossible and should be attempted (11, 17, 47–51). In lieu of isolation, closing the *Dehalobacter* genome(s) may provide more information about variation within the *Dehalobacter* population in SC05 in the future.

Fundamentally, this study contributes to a growing understanding of chloroalkane metabolism, particularly regarding DCM mineralization, but development of a functional assay for DCM assimilation by the Mec cassette is necessary to unequivocally demonstrate this activity. Understanding this culture’s unique metabolism will enable more effective field application and the functional genes presented here are specific biomarkers to survey for successful bioaugmentation.

## Materials and Methods

All chemicals and primers were purchased from Sigma-Aldrich (St. Louis, MO, USA) unless otherwise stated. All gasses were supplied from Linde Canada Inc. (Mississauga, ON, Canada).

### Microbial cultures and growth conditions

SC05-UT is a derivative of SC05, an established enrichment culture used commercially for bioaugmentation (also known as KB-1^®^ Plus CF) originally sampled from a contaminated groundwater site. It dechlorinates CF to DCM and concurrently degrades DCM to carbon dioxide. This culture was maintained in a defined mineral medium without addition of external electron donor for over two years before metagenome sequencing, as previously described (33). DCME is a more recent subculture originally transferred from SC05-UT. This culture was enriched by feeding DCM alone for over 400 days before metagenome sequencing, as previously described (33).

### Metagenome sequencing, assembly, and analysis

These methods are expanded upon in our companion work (35). Briefly, DNA from each subculture was extracted using the Kingfisher Duo Prime MagMax microbiome kit (Thermo Scientific, Waltham, MA). DNA was sequenced by Genome Quebec using both Illumina MiSeq with PCR amplification and PacBio Sequel II technologies. Aliquots of each DNA sample were also used for amplicon sequencing of the V6-V8 region by Genome Quebec as previously described (33). Illumina reads were trimmed using Trimmomatic and FastQC before co-assembly with the PacBio reads using hybridSPAdes (52, 53) (54). Additional quality control, mapping and binning were performed using Anvi’o Snakemake (36, 55, 56). Assembly statistics can be found in Table S6.

Genes were annotated used MetaErg (57); annotations and assigned locus tags for the SC05-UT and DCME metagenomes are shown in supplemental Datasets S1 and S2, respectively. Putative RDase genes were identified by a BLAST search and the presence of the Pfam group PF13486, the hits were filtered for length (>200 amino acids) and possession of at least one iron-sulfur cluster motif (CXXCXXXC), then were each classified with the RDase database (https://rdasedb.biozone.utoronto.ca), using a 90% sequence similarity cutoff to assign each RDase an ortholog group (OG) (19). Each gene in the *Dehalobacterium formicoaceticum mec* cassette was searched for in the metagenome contigs using BLAST to identify homologous proteins (58). Promoters and terminators were predicted using ProPr v2.0 (59).

### Protein extraction

Protein was extracted using an adapted protocol by Murdoch *et al* (23). Briefly, 200 mL of each culture was passed through 0.22 μm Sterivex membrane filter (MilliporeSigma, Burlington, MA) units to capture cells. The filter outlets were then capped with parafilm and 0.5 mL of boiling sodium dodecyl sulfate (SDS) lysis solution (4% SDS in 100 mM Tris/HCl buffer, pH 8.0) was added to each filter unit and incubated at room temperature for one hour on a laboratory shaker. Lysate was extracted from the filter by back pressure using a 3 mL syringe, and filters were rinsed with 0.5 mL of fresh lysis buffer. Lysate mixtures were centrifuged at 21000 xg for 15 min, and supernatant was collected for subsequent trichloroacetic acid precipitation and two acetone washes of the pellet. Protein pellets were air-dried and resuspended in a denaturation buffer (8 M urea, 100 mM Tris– HCl pH 8.0), reduced with 5 mM dithiothreitol (DTT) for 10 min at room temperature, then alkylated with 10 mM iodoacetamide for 10 min at room temperature. Samples were diluted to 4 M urea with water for Bradford Assay quantification (Bio-Rad Laboratories, Inc, Hercules, CA) and trypsin digestion. Protein extracts were stored at -80°C for subsequent high-performance liquid chromatography-tandem mass spectrometry (LC-MS/MS) analysis.

### Metaproteomics

Chromatography was carried out using a PicoTip Emitter (New Objective, Littleton, MA) packed with Reprosil-Pur 120 C18-AQ, 3 μm (Dr. Maisch GmbH, Ammerbuch, Germany). The flow rate was set to 250 nL/min, and the eluents used were 0.1% formic acid in water (A) and 0.1% formic acid in acetonitrile (B). The gradient started at 0% B, followed by a linear gradient to 10% B over 5 min, a linear gradient to 40% B over 48 min, a linear gradient to 95% B over 2 min, a hold at 95% B for 10 min, a linear gradient to 5% B over 2 min, and finally a re-equilibration under the initial conditions of 5% B for 14 min (total runtime 80 min). Liquid samples (5 μL) were injected on a Thermo Scientific Easy nLC 1000 (Thermo Fisher Scientific, Waltham, MA). MS detection was conducted using a Q-Exactive Orbitrap mass spectrometer (Thermo Fisher Scientific, Waltham, MA) equipped with a Nano Electrospray Ionization probe, operating in positive ionization mode, with spray voltage 2.5 kV, capillary temperature 275°C, and s-lens RF level 55. Mass spectra were gathered using a data dependent method as follows; Full MS data was gathered over an m/z range from 400 to 2000 with the mass resolution set to 70k, AGC target 1e6, and a maximum injection time of 30 ms, while data dependant MS2 was gathered using a TOP10 approach, with resolution set to 17.5k, AGC target of 5e4, a maximum injection time of 50 ms, isolation window of 2.0 m/z and NCE 27. Data was processed through X!Tandem/ GPM for initial identification of peptides, and further analyzed through Scaffold. All detected peptides and proteins in SC05-UT and DCME are shown in supplemental information, Dataset S3, and further filtered in Table S3.

### Reductive Dehalogenase Expression and Purification

PCR reagents, restriction enzymes, Gibson Assembly master mix, the 1 kb DNA ladder, and *E. coli* DH5α were purchased from New England Biolabs (Ipswich, MA, USA). The *acdA* gene was PCR-amplified from gDNA extracted from the SC05-UT mixed culture as described above. The TAT sequence was not included in the amplified gene. The primers used for amplification included overhangs complimentary to the *p15TV-L* plasmid backbone as described previously (30, Table S7), such that gene could be inserted by Gibson Assembly into *p15TV-L* (Addgene #26093) linearized by BseRI digestion to produce the expression vector *p15TVL-AcdA*. The plasmid was verified with Sanger sequencing performed by The Centre for Applied Genomics at SickKids Hospital (Toronto, ON, Canada). AcdA was expressed following previously developed methods (42). Briefly, *p15TVL-AcdA* was transformed into *E. coli* BL21(DE3) *cat-araC*-P_BAD_-*suf, ΔiscR::kan, ΔhimA*::Tet^R^, with the *pBAD42-BtuCEDFB* plasmid to co-express the vitamin B_12_ uptake pathway under arabinose control (60, 61). TmrA was expressed using the plasmid *p15TVL-tmrA* with *pBAD42-BtuCEDFB* co-expression in *E. coli* BL21(DE3) *cat-araC*-P_BAD_-*suf, ΔiscR::kan, ΔhimA*::Tet^R^, produced previously (42). The *pBAD42-BtuCEDFB* plasmid was generously provided by the Booker Lab (Pennsylvania State University, PA, USA). *E. coli* BL21(DE3) *cat-araC*-P_BAD_-*suf, ΔiscR::kan, ΔhimA*::Tet^R^, was generously provided by the Antony and Kiley Labs (St. Louis University School of Medicine, MO, USA; University of Wisconsin-Madison, WI, USA).

Large-scale expression of AcdA was performed using 1 L of Luria Broth in a 2 L media bottle. To express AcdA, *E. coli* was first grown aerobically at 37°C, 170 rpm until the OD_600_ reached 0.3-0.5. Expression of the Btu and Suf pathways were induced using a final concentration of 0.2% L-arabinose and the culture was supplemented with 3 μM of hydroxycobalamin hydrochloride. The bottle was sealed with a silicon septum, and the culture was incubated again at 37°C, 170 rpm until the OD_600_ reached 0.6-0.8. The bottle was then put on ice and purged with N_2_ gas for 30 min. After purging, the culture was kept sealed and only handled under an anaerobic atmosphere of N_2_/H_2_/CO_2_ (80:10:10 v/v). Supplements of 50 μM ammonium ferric citrate and 50 μM cysteine were provided and AcdA expression was induced using 0.5 mM Isopropyl β-d-1-thiogalactopyranoside (IPTG). The culture was incubated at 15°C, 170 rpm overnight (18-20 hr). Post-incubation, the culture was pelleted by centrifugation at 4500 xg, 4°C for 20 min. The pellet was stored sealed and at -80°C until it was used for purification.

All lysis and purification steps were done under an anaerobic atmosphere or in a sealed container. In a Coy anaerobic glovebox, the pellet was thawed and resuspended in 10 mL anaerobic lysis buffer [50 mM Tris-HCl pH 7.5, 150 mM NaCl, 0.1% Triton-X100, 5% glycerol, 1 mM tris(2-carboxyethyl)phosphine (TCEP), 50 μg/mL leupeptin, 2 μg/mL aprotinin, 10 mM MgCl_2_], the cells were lysed using BugBuster^®^ protein extraction reagent (MilliporeSigma), 300 μg/mL lysozyme, and 1 μg/mL DNase. The suspended mixture was sealed and incubated outside of the glovebox at room temperature, 70 rpm for 20 min to complete lysis. The crude lysate was clarified by centrifugation at 35 000 xg, 4°C for 20 min. The clarified lysate was transferred into the glovebox and loaded onto a gravity column in with 1 mL nickel-nitrilotriacetic acid resin (Qiagen, Hilden, Germany) that was preequilibrated with wash buffer (50 mM Tris-HCl pH 7.5, 150 mM NaCl, 30 mM imidazole). The protein-bound column was washed with ∼25 mL of wash buffer until protein was no longer detected in the flowthrough by Bradford reagent. AcdA was eluted with 10 mL of elution buffer (50 mM Tris-HCl pH 8, 150 mM NaCl, 300 mM imidazole). The elution was collected and concentrated using a 15 kDa cutoff filter tube (MilliporeSigma) by centrifugation at 3000 xg, 4°C, 30 min. The protein was resuspended in 15 mL of storage buffer (50 mM Tris-HCl pH 7.5, 150 mM NaCl, 1 mM TCEP) and the centrifugation step was repeated. The concentrated protein was stored in liquid N_2_ for long-term stability. The concentration was determined by a Bradford assay and the purity was estimated by SDS-polyacrylamide gel electrophoresis (PAGE) using Image Lab 6.1 software (2022, Bio-Rad Laboratories, Inc.). This expression and purification procedure was repeated for TmrA.

### Chloroalkane Reduction Assay

AcdA dechlorination activity was assessed using an end-point assay. The substrates tested were chloroform; 1,1,1-TCA; 1,1,2-TCA; 1,1-DCA; and DCM. The reactions were performed in 2 mL glass vials with 500 μL inserts and sealed with Teflon lined caps such that there was no headspace. The reaction was carried out in 50 mM Tris-HCl pH 7.5 buffer with 5 mM Ti(III) citrate and 2 mM methyl viologen. A total of 0.31 μg of AcdA or 0.43 μg of TmrA (accounting for purity) was added to the reaction buffer and the respective substrates were suppled to a final concentration of 0.5 mM using saturated water stocks and glass syringes (Hamilton Company, Reno, NV). Once the substate was added, the vial was sealed and incubated upside down at room temperature for 1 hr. Each substrate was tested in triplicate with AcdA, TmrA, and in enzyme free negative controls. The assay was terminated by transferring 400 μL of the reaction into 5.6 mL of acidified (pH < 2) water in a 11 mL headspace autosampler vial immediately sealed with a Teflon coated septum. The samples were analyzed using gas chromatography with flame ionization detection (GC-FID) on an Agilent 7890A GC instrument equipped with an Agilent GS-Q plot column (30 m length, 0.53 mm diameter).

All samples were measured with the same instrument method. The vials were equilibrated to 70°C for 40 min in an Agilent G1888 headspace autosampler. The headspace was sampled, and 3 mL was injected via a heated transfer line and then onto the column by a packed inlet maintained at 200°C. Helium was used as the carrier gas at a flow rate of 11 mL/min. The oven was held at 35°C for 1.5 min, then the temperature ramped up by 15°C/min to 100°C, then the rate was reduced to 5°C/min to 185°C where the temperature was held for 10 min. Finally, the temperature was increased by 20°C/min to 200°C held for 10 min to clear the column. The flame ionization detector was set to a temperature of 250°C. Data were collected and analyzed using Agilent ChemStation Rev. B.04.02 SP1 software. The chromatogram peak areas were translated to liquid concentration using external calibration for each substrate and product. The enzyme activity values were calculated by normalizing the nmol of product measured in the assay by the mass (mg) of purified RDase in the assay and the length of the assay incubation (s). The enzyme free control was reported in nmol product per length of incubation (s).

## Data Availability

Illumina and PacBio metagenome sequencing data and the draft metagenomes are available under GenBank BioProject Accession PRJNA1013980. The assembled metagenomes of SC05-UT and DCME are available at NCBI under Genbank accession numbers JAWDGN00000000 and [JAWDGO00000000](https://www.ncbi.nlm.nih.gov/nuccore/JAWDG000000000.1/), respectively. The versions used here are the first versions, JAWDGN010000000 and JAWDGO010000000. Illumina paired reads have been deposited in the Sequence Read Archive under accession numbers SRR27458442 (SC05-UT) and SRR27458441 (DCME). PacBio reads were deposited in the SRA under accession numbers SRR25941764 (SC05-UT) and SRR25941763 (DCME). Fasta files of all draft MAGs in the DCME and SC05-UT metagenomes are available at https://figshare.com/s/784a246f0cb57c3cca90 and https://figshare.com/s/74a665aec10a8bc9622f, respectively.

## Acknowledgments

This work was funded by the Natural Science and Engineering Research Council (NSERC) though a Discovery Grant to E.A.E. and Doctoral Scholarships to O.B., by an Ontario Graduate Scholarship to K.P., as well as by a Genome Canada Bioinformatics and Computational Biology (BCB project 285MPR) subgrant to E.A.E. and R.M. Authors acknowledge funding from the Canada Research Chairs Program to R.M. and E.A.E. The funders had no role in study design, data collection and interpretation, or the decision to submit the work for publication. We thank SiREM for providing us with the culture studied in this work and Elizabeth Phillips for her valuable insights into caring for the culture. We acknowledge Rob Flick for performing the mass spectrometry and Miya Tseng-West for assistance in cloning. We would also like to thank the Booker Lab from Pennsylvania State University for gifting the *pBAD42-BtuCEDFB* plasmid, and the Antony Lab from the St. Louis University School of Medicine and Kiley lab from the University of Wisconsin-Madison for gifting us the *E. coli* BL21(DE3) *cat-araC*-P_BAD_-*suf, ΔiscR::kan, ΔhimA*::Tet^R^ strain.

## Contributor Roles

E.A.E., R.M, O.B., and K.P. conceived of the project. O.B. established the microbial cultures, performed metagenomic sequencing and assembly and proteomic analyses. K.P. performed heterologous expression and enzyme assays. The manuscript was written by O.B., K.P. and E.A.E. All authors contributed to manuscript revision and have approved the submitted version.

## Notes

### Competing Interest Statement

The authors have declared no competing interest.

### Summary of Updates

The introduction has been expanded to further clarify the results, and the results section has been amended with a more in-depth discussion of two subcultures of SC05, highlighting a key genomic neighbourhood important to the active *Dehalobacter* in these cultures.

