## Supplementary Information for "From *mec* cassette to *rdhA*: a key *Dehalobacter* genomic neighborhood in a chloroform and dichloromethane–transforming microbial consortium"

#### SUPPLEMENTARY DATASETS

All datasets are deposited in FigShare as follows:

**Dataset S1:** Annotation of the metagenome of the microbial consortium SC05-UT.
https://doi.org/10.6084/m9.figshare.25145888.v1
**Dataset S2:** Annotation of the metagenome of the microbial consortium DCME.
https://doi.org/10.6084/m9.figshare.25146014.v1
**Dataset S3:** Proteomic statistical analysis—all values from LC-MS/MS read analysis including X!Tandem algorithm intensity score and identification of peptides (S3A) and proteins (S3B)
https://doi.org/10.6084/m9.figshare.25146029.v1
**Dataset S4:** All raw data and relevant information for the AcdA dechlorination assays
https://doi.org/10.6084/m9.figshare.25146044.v1

#### SUPPLEMENTARY TEXTS

**Text S1:** Ortholog group 97 …………………………...S4

**Text S2:** Assembly statistics ……………..………….S5

**Text S3:** Metagenomic sequencing of SC05-UT and DCME …………………………...S6

**Text S4:** Identification *acd* and *mec* cassette operon …………………………...S7

**Text S5:** 1,1,1-TCA and 1,1-DCA dechlorination by SC05-UT …………………………...S8

**Text S6:** Putative DCME mec contig quality ……………………….S9

**Text S7:** Heterologous expression of AcdA ……………………….S14

#### SUPPLEMENTARY TABLES

**Table S1:** Metagenome bins and quality statistics …...Accompanying Excel

**Table S2:** List of *rdhA* in SC05-UT metagenome …...Accompanying Excel

**Table S3:** Protein hits from SC05-UT and DCME *Dehalobacter* …...Accompanying Excel

**Table S4:** Mec cassette and WLP proteomic expression …...Accompanying Excel

**Table S5:** Metagenome assembly statistics ………………………S5

**Table S6:** Promoters and terminators in *acdA*/ *mec* region ……………………….S7

**Table S7:** AcdA cloning primers …..…………………S16

**Table S8:** Expression plasmids …..…………………S16

#### SUPPLEMENTARY FIGURES

**Figure S1:** Similarity matrix of OG 97 members ……………………….S4

**Figure S2:** Community compositions of metagenome samples ……………………….S6

**Figure S3:** *mec* cassette and *acdA* operons in SC05-UT ……………………….S7

**Figure S4:** SC05-UT dechlorination of 1,1,1-TCA ……………………….S8

**Figure S5:** Quality analysis of putative DCME *mec* contig ……………………...S10

**Figure S6:** Read mapping of DCME and SC05-UT contig ……………………...S12

**Figure S7:** Alignment of DCME/SC05-UT gene neighborhoods ……………………...S13

**Figure S8:** SDS-PAGE of AcdA ……………………...S17

#### TEXT S1. Ortholog group 97

Ortholog groups (OGs) are comprised of reductive dehalogenases (RDases) that have over 90% sequence identity to one another (1). Here we show the classified members of OG 97 and the similarity of their amino acid sequences to confirm placement of AcdA in the group and to show the most similar sequences to AcdA (Figure S1).

| CfrA | CtrA | ThmA | DcrA | RdhA D8M | TmrA | AcdA | DCME AcdA | |
|---|---|---|---|---|---|---|---|---|
| | 93.9% | 95.4% | 95.0% | 94.5% | 95.4% | 95.8% | 95.2% | CfrA (*Dehalobacter* sp. CF) |
| 28 | | 97.8% | 93.9% | 93.2% | 94.1% | 93.7% | 93.2% | CtrA (*Desulfitobacterium* sp. PR) |
| 21 | 10 | | 95.6% | 95.0% | 95.8% | 95.4% | 95.0% | ThmA (*Dehalobacter* sp. THM1) |
| 23 | 28 | 20 | | 94.1% | 95.2% | 96.7% | 96.7% | DcrA (*Dehalobacter* sp. DCA) |
| 25 | 31 | 23 | 27 | | 96.9% | 96.3% | 96.1% | RdhA D8M_v2_40029 (*Dehalobacter* sp. 8M) |
| 21 | 27 | 19 | 22 | 14 | | 96.7% | 96.1% | TmrA (*Dehalobacter* sp. UNSWDHB) |
| 19 | 29 | 21 | 15 | 17 | 15 | | 99.1% | AcdA (SC05-UT *Dehalobacter*) |
| 22 | 31 | 23 | 15 | 18 | 18 | 4 | | AcdA (DCME *Dehalobacter*) |

**FIGURE S1** Similarity matrix of all characterized OG97 RDases showing percent amino acid identity (green, upper right) and number of residue differences (blue, lower left). Darker shades indicate higher similarity. An amino acid alignment was built using the Geneious v8.1.9 MUSCLE plugin. Protein accession numbers for sequences are as follows: CfrA, AFV05253 (2); CtrA, AGO27983 (3); ThmA, ANI21407 (4); DcrA, AFV02209 (2); RdhA D8M_v2_40029, No accession available (5); TmrA, WP_034377773 (6); AcdA, locus tag JAWDGN_38927 (7); DCME AcdA, locus tag JAWDGO_25608. Figure adapted from (7).

#### TEXT S2. Assembly statistics

This section contains the assembly statistics of SC05-UT and DCME metagenomes (Table S5). Other metagenomic and proteomic data is in the accompanying excel file, as listed in the Table of Contents, including metagenomic *rdhA* identification in SC05-UT (Table S2), genomic and proteomic identification of the Mec cassette and Wood-Ljungdahl pathway proteins in SC05-UT and DCME *Dehalobacter* MAGs (Table S1), and DAS Tool binning statistics from both metagenomes (Table S3).

**TABLE S5.** Assembly statistics for the metagenomes of each sampled culture.

| | **DCME** | **SC05-UT** |
|---|---|---|
| Total nucleotides (Mb) | 243.5 | 413.5 |
| Number of contigs | 200,127 | 467,998 |
| Num Contigs > 100 kb | 233 | 365 |
| Num Contigs > 50 kb | 561 | 788 |
| Num Contigs > 20 kb | 1,692 | 1,803 |
| Num Contigs > 10 kb | 3,504 | 3,987 |
| Num Contigs > 5 kb | 6,087 | 7,597 |
| Num Contigs > 2.5 kb | 9,306 | 14,297 |
| Longest Contig | 1,675,588 | 1,741,616 |
| Shortest Contig | 56 | 56 |
| L50 | 2,232 | 8,122 |
| N50 | 15,727 | 4,648 |
| Number of DAS Tool bins (see Table S1) | 40 | 51 |

#### TEXT S3. Metagenomic sequencing of SC05-UT and DCME

Below, we show the community compositions of each DNA sample sent for metagenomic sequencing. Two separate DNA samples were taken for each culture, one sent for paired-end sequencing on Illumina MiSeq and one sent for long-read sequencing on a PacBio Sequel II. To ensure community compositions of each sample were similar, a subsample of each DNA sample was sent for 16S amplicon sequencing. These results are compared in Figure S2. Relative abundances of microorganisms cannot be calculated from the metagenomic reads directly, because of PCR bias during an amplification step prior to shotgun sequencing.

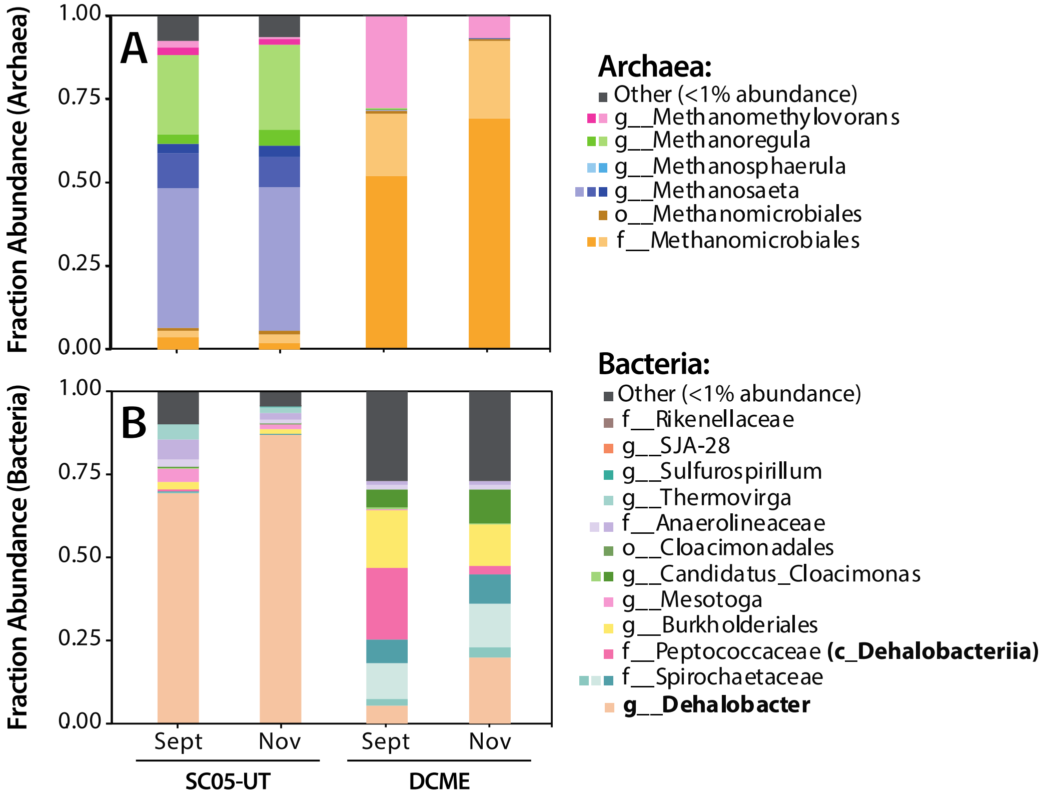


**FIGURE S2.** Community compositions determined by 16S amplicon sequencing of **A**) archaea and **B**) bacteria in each DNA sample sent for metagenome sequencing (either Illumina in September or PacBio in November 2021). Each colour represents one ASV, as classified by SILVA. Organisms of interest are bolded.

#### TEXT S4. Identification *acdABC* and *mec* cassette operon

Here we show a depiction of a contig subsection containing the SC05-UT *mec* cassette and *acdABC* including strong promoters and terminators predicted in this region by ProPr v2.0 (Figure S3, Table S6) (8). There are several strong promoters predicted in this region, including immediately upstream of *acdA*, but notably there is no promoter predicted near *mecA* (Figure S3). It is possible the terminator between the nearest promotor and *mecA* is not very strong, allowing *mec* cassette expression, or perhaps a promoter exists that has remained undetected by our methods.

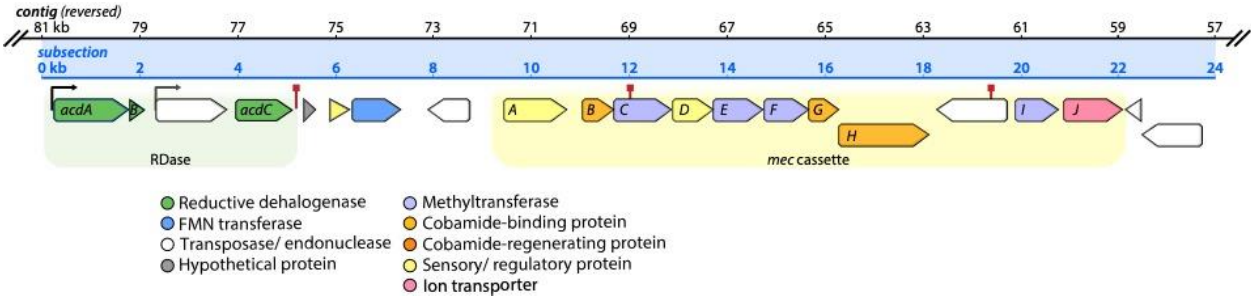


**FIGURE S3.** Position of *mec* cassette and *acdABC* operons on SC05-UT *Dehalobacter* contig (JAWDGN010000066.1). A 24 kb subsection of the contig is shown in blue, extracted from the 81-57 kb range of the reversed contig, labelled in black to show contig location. The *mec* cassette is highlighted in yellow and the RDase is highlighted in green. Other genes are colored by function. Predicted promoters and terminators on the positive strand are marked with black arrows and red squares, respectively (described in Table S6: promoter_1, promoter_2, terminator_1, terminator_2, terminator_3).

**TABLE S6.** Strong promoters and terminators predicted on contig subsection shown in Figure S3. dG: Gibb's free energy of terminator stem-loop formation, TSS: transcription start site.

| ID | Start position (kb) | End position (kb) | Strand | dG (kcal/ mol) | Promoter Quality Score | Predicted TSS (kb) |
|---|---|---|---|---|---|---|
| promoter_1 | 80835 | 80764 | + | NA | 0.98 | 80759 |
| promoter_2 | 78778 | 78707 | + | NA | 0.93 | 78707 |
| promoter_3 | 61227 | 61156 | - | NA | 0.95 | 61228 |
| promoter_4 | 70082 | 70011 | - | NA | 0.93 | 70081 |
| promoter_5 | 72019 | 71948 | - | NA | 0.95 | 72019 |
| terminator_1 | 77136 | 77121 | + | -13.4 | NA | NA |
| terminator_2 | 70438 | 70422 | + | -11.3 | NA | NA |
| terminator_3 | 63020 | 63004 | + | -10.4 | NA | NA |

#### TEXT S5. 1,1,1-TCA and 1,1-DCA dechlorination by SC05-UT

To determine if the SC05-UT culture can dechlorinate 1,1,1-trichloroethane (TCA) and 1,1-dichloroethane (DCA) *in vivo*, 20 mL aliquots of SC05-UT were added to 80 mL of medium each in each of four 160 mL serum bottles and sealed with blue butyl rubber stoppers. The inoculations were performed in an anaerobic chamber (Coy) supplied with a $N_2/H_2/CO_2$ gas mix (80:10:10 v/v) and were purged with $N_2/CO_2$ (80:20 v/v) for 30 min before feeding. A glass syringe was used to feed 7 µL (0.5 mM aqueous) 1,1,1-TCA to each replicate. Each culture was amended with electron donor at a 1:1 electron equivalent ratio to 1,1,1-TCA, as follows. Two replicates received 6.6 mL $H_2/CO_2$ (20:80 v/v), and two replicates received 140 µL of 400 mM formate. A second amendment of electron donor was performed on day 138. The same degradation pattern was seen in both donor scenarios (Figure S4).

Bottles were measured periodically by gas chromatography, wherein 1,1,1-TCA, 1,1-DCA, CA, and methane were quantified. Each 0.3 mL headspace sample was injected into a Hewlett-Packard 5890 Series II gas chromatograph (GC) fitted with a GSQ column (30-m-by-0.53-mm [inner diameter] PLOT column; J&W Scientific, Folsom, CA). The GC carrier gas pressure was set initially to 100 kPa, and the oven temperature was programmed to hold at 50°C for 1.5 min, then increase to 180°C at 60°C/min and hold for 2.5 min. Calibration was performed using external standards. Results are shown in Figure S4.

Though the dechlorination of 1,1,1-TCA occurs at rate approximately 10-fold slower than the dechlorination of CF by the parent culture, it might be sped up after several re-feedings and more frequent electron donor amendment.

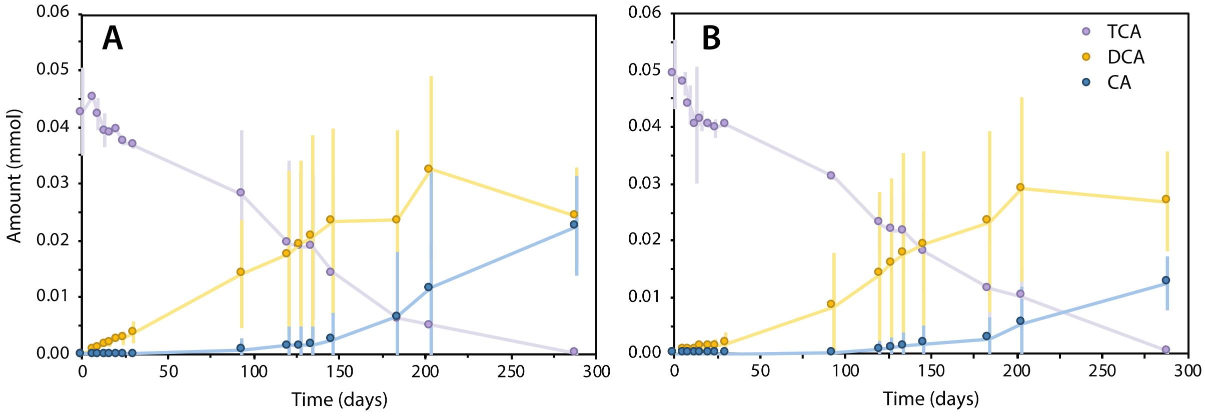


**FIGURE S4.** Dechlorination of 1,1,1-TCA and 1,1-DCA in SC05-UT using A) formate or B) hydrogen as an electron donor over time. (n=2, error bars indicate range).

#### TEXT S6. Putative DCME *mec* contig quality

Here we describe the consideration of a putative end to the *mec* cassette in the DCME *Dehalobacter* MAG, which was identified through BLAST to encode *mecGHIJ* on the end of a 53.8 kb contig assigned to an Anaerovoracaceae bin (contig JAWDGO010000514) in the DCME metagenome (Figure S5). Due to PCR amplification during the preparation of our metagenomic reads, coverage of contigs in both the SC05-UT and DCME metagenome assemblies is uneven and mountainous, especially for low abundance organisms. Therefore, we cannot rely on uneven coverage alone to discard any contig from a bin, and must look closer to examine the relevance and taxonomy of this potential *mec* cassette.

Contig JAWDGO010000514 has short-read coverage with a mean of 17.2—less than half the mean coverage of the DCME *Dehalobacter* bin (42.3)—and several coverage gaps (Figure S5A, highlighted in orange). The *mecGHIJ* region has higher short-read coverage than the rest of the contig, but no PacBio HiFi long reads mapped to *mecGH,* or connected the *mec* cassette to the rest of the contig (Figure S5B, regions with no long-read coverage highlighted in red). This uneven short read coverage, as well as the tenuous connection of the *mec* cassette to the rest of the contig, suggests a potential misassembly between these two regions. Furthermore, the contig's GC content (47.1%) was higher than the rest of the neighborhood in both DCME (40.8%) and SC05-UT (40.3%).

Looking downstream of the no-coverage region, when compared to the *mec* cassette neighbourhood in the SC05-UT *Dehalobacter*, the only homologous genes are a DNA-binding protein and a recombinase (Figure S5C). These are present in many bacterial genomes as a by-product of mobile elements, and do not imply that this contig is linked to *Dehalobacter*. For the reasons above, this contig was considered unreliable—perhaps a faulty assembly, and very likely not the end of the *Dehalobacter* cassette.

Nonetheless, this contig's *mecGH* sequences are divergent from those seen on the *Dehalobacter* contigs, and may represent an additional copy of the *mec* cassette encoded by a low-abundance organism (whether misassembled to the Anaerovoracaceae contig or not). Thus, the contig was still used for proteomic searching. These proteins were not detected in the proteome

(Supplementary Dataset S3), suggesting that this organism is not a major player in DCM degradation in the culture.

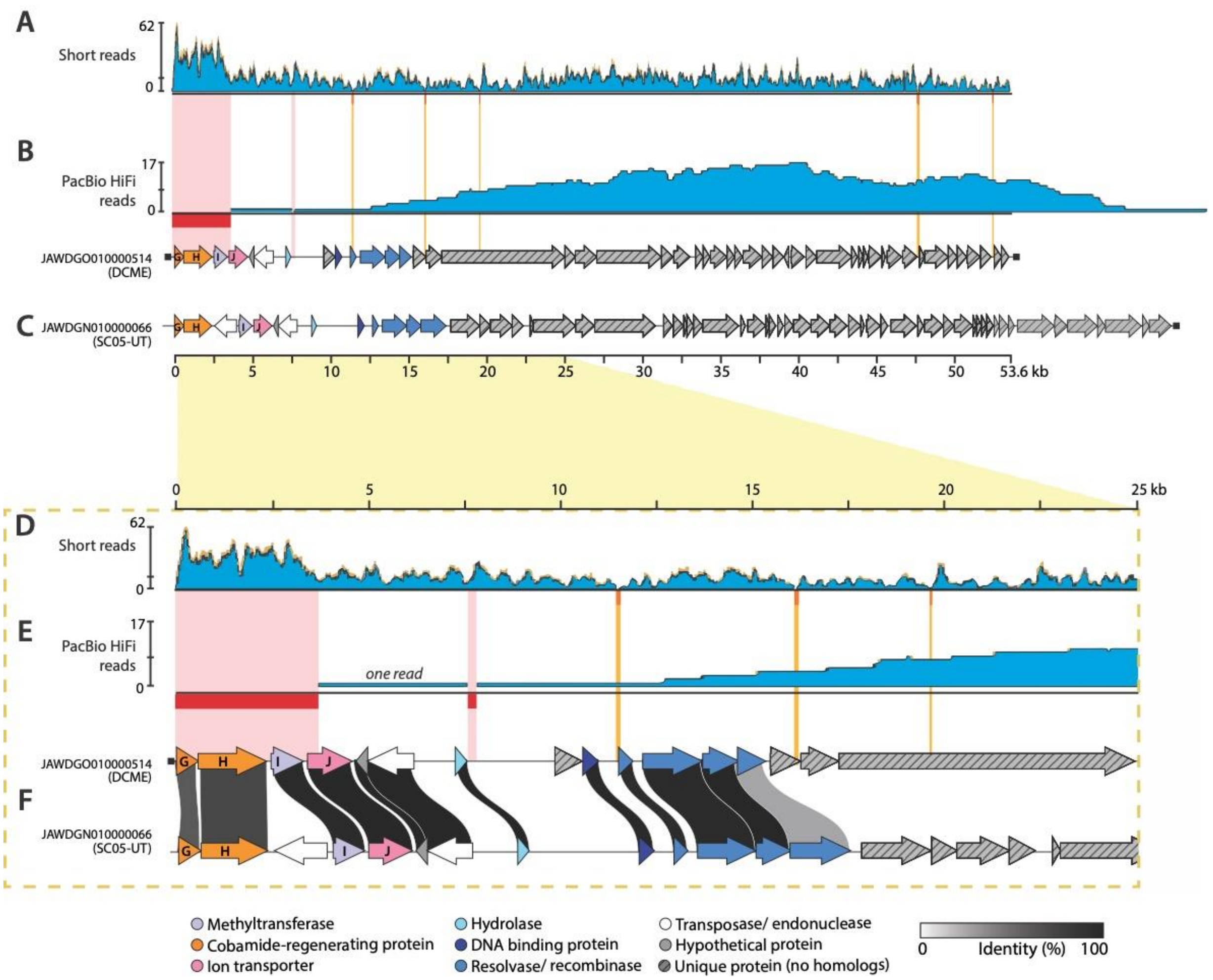


**FIGURE S5.** Quality analysis of putative *mec* contig (JAWDGO010000514) from the DCME metagenome. Panels A-C show reads mapped to the full contig, and panels D-F zoom on the *mec* fragment (highlighted in yellow). **A, D**) Short read mapping to contig, with zero-coverage regions highlighted in orange. **B, E**) Long read mapping of PacBio HiFi reads to contig, with zero-coverage regions highlighted in red. **C, F**) Homology of putative contig to analogous SC05-UT contig (JAWDGN010000066, downstream of *mecG*). Homologous genes are color-coded by function; links between homologs are colored by percent nucleotide identity. Genes with no local homologs are grey striped. Ends of a contig are capped with black squares.

The SC05-UT *Dehalobacter* contig JAWDGN010000066—containing the *mec* cassette and *acdA*—has high long and short-read coverage without any gaps, which provides confidence in our *Dehalobacter* assembly (Figure S6B). Due to the relatively poor quality of the DCME

*Dehalobacter* MAG and low coverage overall compared to the SC05-UT MAG (mean: 37.7 vs. 1027.7), read mapping to contig JAWDGO010000051 (encoding *mecABCDEFGH*) also resulted in some gaps (Figure S6A). One gap in the HiFi read coverage exists in the *acd-mec* neighborhood (Figure S6C). Luckily, this region is covered by short reads in the DCME metagenome, and many HiFi reads in the SC05-UT assembly. Some "wavy" coverage is seen in both *Dehalobacter* contigs due to PCR bias from an amplification step prior to shotgun sequencing, and higher coverage exists over some repeated elements like transposases. Population variation may also contribute to uneven coverage, which may be resolved with genome closure after further sequencing.

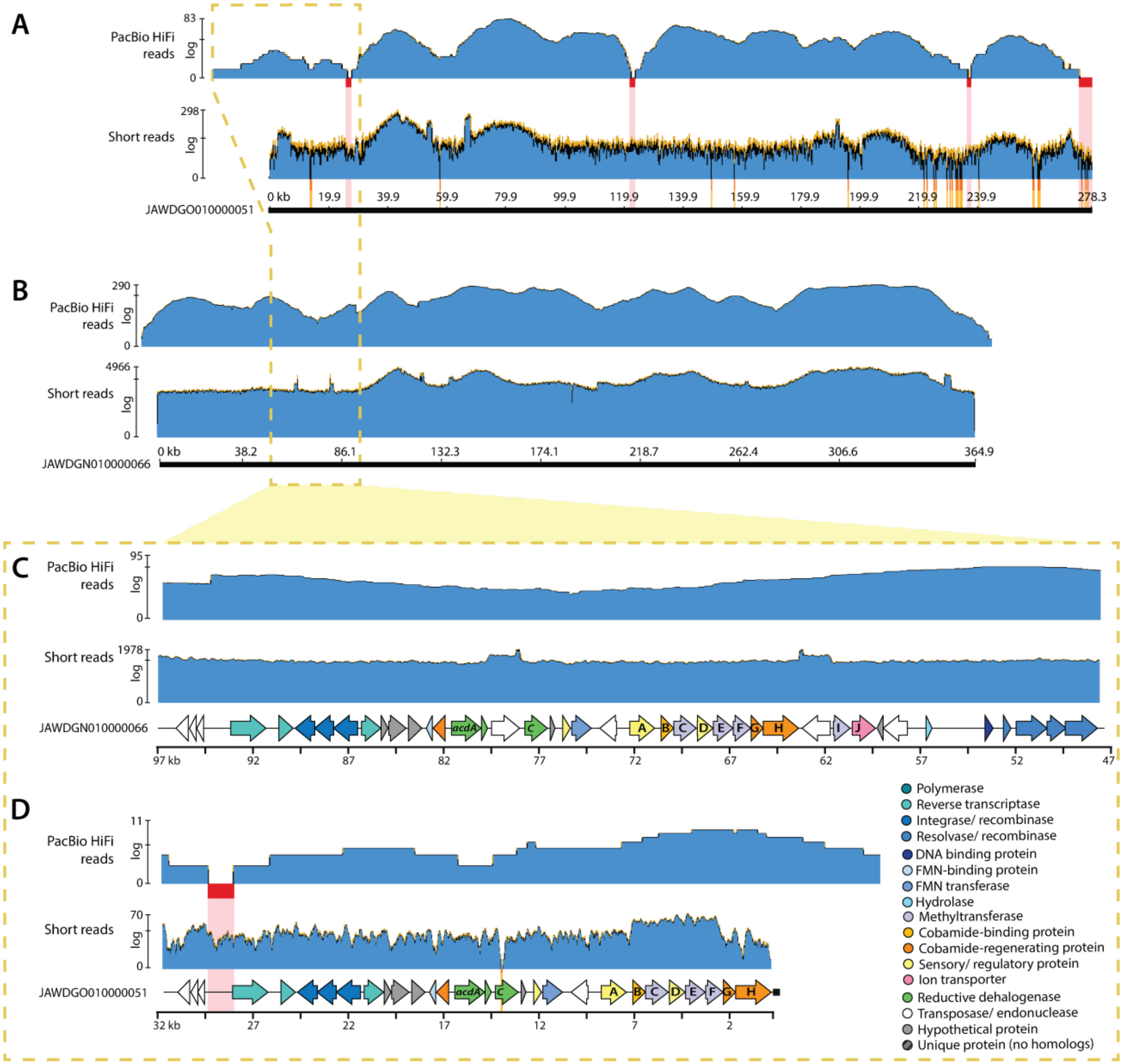


**FIGURE S6.** Read mapping of Illumina paired-end short reads and PacBio HiFi long reads to *mec* contigs **A)** DCME metagenome: contig JAWDGO010000051 **B**) SC05-UT metagenome: contig JAWDGN010000066. Panels C and D zoom on the *acd-mec* neighborhood. Zero-coverage regions in short read mapping are highlighted in orange; with long read zero-coverage regions highlighted in red. Gene neighborhood containing *mec* cassette and *acdABC* highlighted in green.

After long-read mapping to the DCME contig JAWDGO010000051, the mapped PacBio HiFi reads overhang the downstream end of the *mec* cassette (Figure S6A, S6C). These reads were used to perform contig extension, and correct ambiguous bases in the paired-end read mapping, revealing *mecIJ* following a transposase, syntenic to the SC05-UT contig JAWDGN010000066. An alignment of these sequences, including the extension predicted from HiFi reads, is shown in

Figure S7. With this contig extension, the end of the DCME *Dehalobacter* MAG *mec* cassette mirrors that of SC05-UT *Dehalobacter* MAG.

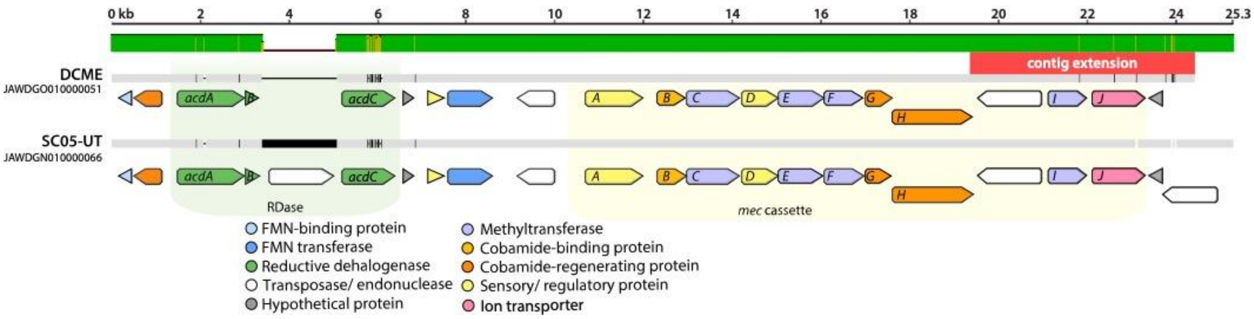


**FIGURE S7.** Alignment of RDase-*mec* gene neighborhoods in DCME and SC05-UT until end of contig JAWDGO010000051. Region amended via contig extension highlighted in red. Local percent identity (sliding window = 15) is shown in green, with disagreements in yellow. Identical sequences are grey, and differences are highlighted in black. Genes are colored by function.

### TEXT S7. Heterologous expression of AcdA

This section contains the nucleotide and amino acid sequences of AcdA used for heterologous expression, an alignment of the protein sequences from DCME and SC05-UT, as well as primers used for amplification of the gene from metagenomic DNA (Table S7), expression plasmids used (Table S8), and SDS-PAGE gel visualization of expression and purification of AcdA (Figure S5). Raw enzyme assay data is shown in the accompanying excel file (Dataset S4). Only the AcdA from SC05-UT was expressed for activity assays since this culture was actively dechlorinating CF.

> *acdA* | SC05-UT *Dehalobacter* nucleotide sequence

```
1     atggacaaggaaaaaagtaa caacgataagccggcaacaa aaattaatcgcagacaattc cttaaatttggagctggagc ttcttcgggtattgcaattg
101   ccactgcagctactgcattg ggagggaaatcacttatcga tcccaaacaggtatatgctg gaacggtcaaggaactggat gaacttccctttaatatccc
201   ggcagactacaaaccgttta ccaatcaaaggaatatattt ggccaggctgtattgggagt acccgaacctctagcacttc aagagcgttttgatgaagta
301   agatggaatggttggcagac agatggttcgcccggtctta ctgtacttgatggtgcggct gctcgtgcaagctttgccgt tgattattatcttaacgggg
401   aaaatagcgcctgcagggcc aataaaggtttttttgaatg gcatcccaaagtgcccgagc tgaactttaagtggggcgat ccggagagaaatattcattc
501   ccccggtgtaaaaagtgccg aagaaggaacgatggcagta aaaaaaatagctagattttt cggcgctgctaaagctggga tagcgccttttgacaaacgt
601   tgggtttttactgaaacggc tgcctttgttaaaacgcctg agggtgaaagtctgaaattt atccctccggattttgggtt tgagcccaagcatgtaatct
701   cgatgattatcccacagtcg ctagaaggaataaagtgtgc cccgtcctttttaggatcag ctgaatatggattaagttat gcccagattggatatgctgc
801   attcggtttatccatgttta ttaaagatctgggatatcat gcggttccaatcggagctga cagtgcattagctgtaccta tagctattcaggcgggtctg
901   ggggaatacagcaggtcggg gctaatgattacgcctgaat ttggttcaaatgttagactc tgtgaagtatttactgacat gcctttaaatcatgataaac
1001  ctatttcattcggagtaact gaattttgcaaaacctgcaa aaaatgcgctgaagcatgcc cccctcaagctattagctat gaagatcctaccattgatgg
1101  acctcgtgggcaaatgcata attcgggaataaagagatgg tatgttgacccggtgaagtg ctttgaattctggtcgcgtg ataacgtcagaaactgctgc
1201  ggagcttgtatagctgcttg cccatttactaagccggaag cctggcaccataccttaatt aggagtctagtaggagcacc tgttattactccattcatga
1301  aagatgtggatgatattttt ggatacggaaagccgaatga tgaaaaagcgatagcagatt ggtggaaataa
```

> AcdA | SC05-UT *Dehalobacter* Amino acid sequence, TAT sequence is bolded and underlined

**MDKEKSNNDKPATKINRRQFLKFGAGASSGIAIATAATALGGKSL**IDPKQVYAGTVKELDELPF
NIPADYKPFTNQRNIFGQAVLGVPEPLALQERFDEVRWNGWQTDGSPGLTVLDGAAARASFAVD
YYLNGENSACRANKGFFEWHPKVPELNFKWGDPERNIHSPGVKSAEEGTMAVKKIARFFGAAKA
GIAPFDKRWVFTETAAFVKTPEGESLKFIPPDFGFEPKHVISMIIPQSLEGIKCAPSFLGSAEY
GLSYAQIGYAAFGLSMFIKDLGYHAVPIGADSALAVPIAIQAGLGEYSRSGLMITPEFGSNVRL
CEVFTDMPLNHDKPISFGVTEFCKTCKKCAEACPPQAISYEDPTIDGPRGQMHNSGIKRWYVDP
VKCFEFWSRDNVRNCCGACIAACPFTKPEAWHHTLIRSLVGAPVITPFMKDVDDIFGYGKPNDE
KAIADWWK

> Protein alignment of DCME *Dehalobacter* AcdA, SC05-UT *Dehalobacter* AcdA, differences are highlighted in red and the consensus sequence is in the center. Positive matches denote different amino acids that have similar chemical properties.

Identities = 452/456 (99%),
Positives = 453/456 (99%), Gaps = 0/456 (0%)

```
DCME AcdA     1   MDKEKSNNDKPATKINRRQFLKFGAGASSGIAIATAATALGGKSLIDPKQVYAGTVKELD
                  MDKEKSNNDKPATKINRRQFLKFGAGASSGIAIATAATALGGKSLIDPKQVYAGTVKELD
SC05-UT AcdA  1   MDKEKSNNDKPATKINRRQFLKFGAGASSGIAIATAATALGGKSLIDPKQVYAGTVKELD

DCME AcdA     61  ELPFNIPADYKPFTNQRNIFGQAVLGVPEPLALEERFAEVRWNGWQTDGSPGLTVLDGAA
                  ELPFNIPADYKPFTNQRNIFGQAVLGVPEPLAL+ERF EVRWNGWQTDGSPGLTVLDGAA
SC05-UT AcdA  61  ELPFNIPADYKPFTNQRNIFGQAVLGVPEPLALQERFDEVRWNGWQTDGSPGLTVLDGAA

DCME AcdA     121 ARASFAVDYYLNGENSACRANKGFFEWHPKVPELNFWWGDPERNIHSPGVKSAEEGTMAV
                  ARASFAVDYYLNGENSACRANKGFFEWHPKVPELNF WGDPERNIHSPGVKSAEEGTMAV
SC05-UT AcdA  121 ARASFAVDYYLNGENSACRANKGFFEWHPKVPELNFKWGDPERNIHSPGVKSAEEGTMAV

DCME AcdA     181 KKIARFFGAAKAGIAPFDKRWVFTETAAFVKTPEGESLKFIPPDFGFEPKHVISMIIPQS
                  KKIARFFGAAKAGIAPFDKRWVFTETAAFVKTPEGESLKFIPPDFGFEPKHVISMIIPQS
SC05-UT AcdA  181 KKIARFFGAAKAGIAPFDKRWVFTETAAFVKTPEGESLKFIPPDFGFEPKHVISMIIPQS

DCME AcdA     241 LEGIKCAPSFLGSAEYGLSYAQIGYAAFGLSMFIKDLGYHAVPIGADSALAVPIAIQAGL
                  LEGIKCAPSFLGSAEYGLSYAQIGYAAFGLSMFIKDLGYHAVPIGADSALAVPIAIQAGL
SC05-UT AcdA  241 LEGIKCAPSFLGSAEYGLSYAQIGYAAFGLSMFIKDLGYHAVPIGADSALAVPIAIQAGL

DCME AcdA     301 GEYSRSGLMITPEFGSNVRLCEVFTDMPLNHDKPISFGVTEFCKTCKKCAEACPPQAISY
                  GEYSRSGLMITPEFGSNVRLCEVFTDMPLNHDKPISFGVTEFCKTCKKCAEACPPQAISY
SC05-UT AcdA  301 GEYSRSGLMITPEFGSNVRLCEVFTDMPLNHDKPISFGVTEFCKTCKKCAEACPPQAISY

DCME AcdA     361 EDPTIDGPRGQMHNSGIKRWYVDPVKCFEFWSRDNVRNCCGACIAACPFTKPEAWHHTLT
                  EDPTIDGPRGQMHNSGIKRWYVDPVKCFEFWSRDNVRNCCGACIAACPFTKPEAWHHTL
SC05-UT AcdA  361 EDPTIDGPRGQMHNSGIKRWYVDPVKCFEFWSRDNVRNCCGACIAACPFTKPEAWHHTLI

DCME AcdA     421 RSLVGAPVITPFMKDVDDIFGYGKPNDEKAIADWWK
                  RSLVGAPVITPFMKDVDDIFGYGKPNDEKAIADWWK
SC05-UT AcdA  421 RSLVGAPVITPFMKDVDDIFGYGKPNDEKAIADWWK
```

**TABLE S7.** Primers used for amplification of *rdhA* genes for cloning. Bolded sequences are complimentary to the *rdhA* gene, non-bolded is an extension for Gibson assembly. Primers are from Picott et al. 2022, the primer name corresponds to the previously published primer name.

| **Primer Name (from Picott et al. 2022)** | **Sequence (5'→3')** | **Purpose** |
|---|---|---|
| *OG97_no_TAT_F* | TTGTATTTCCAGGGC**ATGATCGATCCCAAACAGGTA** | Amplify AcdA without TAT signal peptide sequence for Gibson Assembly into *p15TV-L* |
| *CfrA_DcrA_R* | CAAGCTTCGTCATCA**TTATTTCCACCAATCTGCTATC** | Reverse primer to amplify AcdA for Gibson Assembly into *p15TV-L*. |

**TABLE S8.** Expression plasmids used in this work.

| **Plasmid** | **Encoded Enzyme** | **Tags** | **Induction** | **Source** |
|---|---|---|---|---|
| *p15TVL-AcdA* | AcdA (no TAT) | N-term 6xHis | IPTG | This work |
| *P15TVL-TmrA* | TmrA (no TAT) | N-term 6xHis | IPTG | (9) |
| *pBAD42-BtuCEDFB* | BtuB, BtuC, BtuD, BtuE, BtuF | N/A | Arabinose | Booker lab (Pennsylvania State University) (10) |

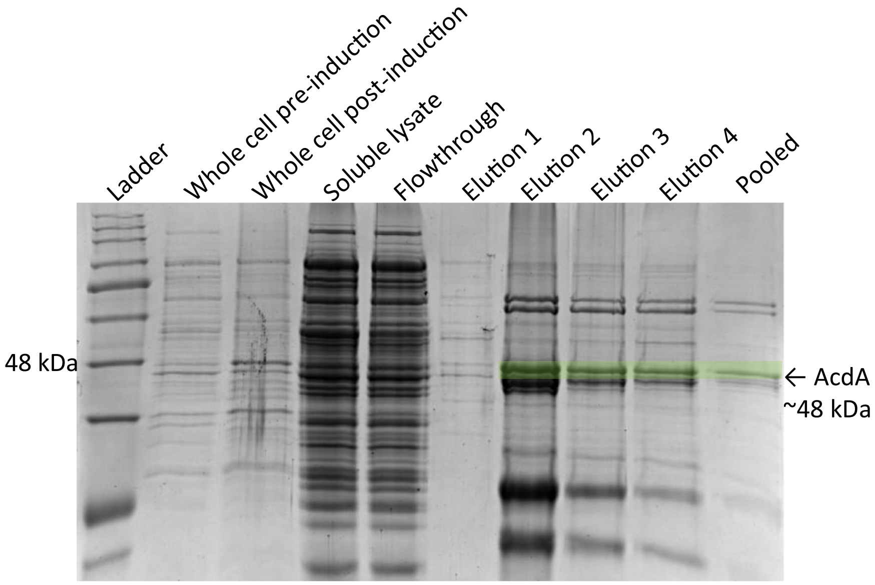


**FIGURE S8.** SDS-PAGE of AcdA (47.9 kDa) expression and purification by nickel affinity chromatography. The whole cell lane was taken pre-induction and 18 hrs post-induction, the cell pellet was lysed using BugBuster extraction reagent (Millipore) and the soluble fraction was imaged. The protein was purified by gravity chromatography using 1 mL of Ni-NTA resin from Qiagen, the flowthrough was collected for imaging. The elution fractions were combined and concentrated by a 30 kDa cut-off Millipore filter tube (Pooled). The ladder is Frogga Bio BLUeye pre-stained protein ladder.
